# One-Step Rapid Quantification of SARS-CoV-2 Virus Particles via Low-Cost Nanoplasmonic Sensors in Generic Microplate Reader and Point-of-Care Device

**DOI:** 10.1101/2020.06.09.142372

**Authors:** Liping Huang, Longfei Ding, Jun Zhou, Shuiliang Chen, Fang Chen, Chen Zhao, Yiyi Zhang, Jianqing Xu, Wenjun Hu, Jiansong Ji, Hao Xu, Gang L. Liu

## Abstract

The spread of SARS-CoV-2 virus in the ongoing global pandemics has led to infections of millions of people and losses of many lives. The rapid, accurate and convenient SARS-CoV-2 virus detection is crucial for controlling and stopping the pandemics. Diagnosis of patients in the early stage infection are so far limited to viral nucleic acid or antigen detection in human nasopharyngeal swab or saliva samples. Here we developed a method for rapid and direct optical measurement of SARS-CoV-2 virus particles in one step nearly without any sample preparation using a spike protein specific nanoplasmonic resonance sensor. We demonstrate that we can detect as few as 30 virus particles in one step within 15 minutes and can quantify the virus concentration linearly in the range of 10^3^ vp/ml to 10^6^ vp/ml. Measurements shown on both generic microplate reader and a handheld smartphone connected device suggest that our low-cost and rapid detection method may be adopted quickly under both regular clinical environment and resource-limited settings.

## Introduction

The ongoing COVID-19 pandemic is caused by the respiratory infections of SARS-CoV-2 (Severe Acute Respiratory Syndrome Coronavirus 2) virus between humans^1, 2^. It is evident that the infection may be effectively reduced and eventually controlled with swift and large-scale testing for early diagnostics^3-6^. Current testing methods include PCR nucleic acid detection with either thermal cycling^7, 8^ or isothermal amplification^9-11^, serological IgM/IgG antibody testing^12^ as well as viral protein detections^13^ in nasopharyngeal swap. The PCR based nucleic acid testing is very sensitive and accurate in early infection diagnostics but it generally requires multiple and lengthy processes including virus lysis, RNA extraction, reverse transcription and amplification and is prone to sample contamination^14, 15^. Serological IgM/IgG antibody testing is a very rapid and potentially point-of-care detection method for epidemiology studies of past viral infections, although not adequate for early diagnosis, since the serological antibody presence will not occur until a couple of weeks after the initial viral infection^16-18^. There were some reported and commercially available viral antigen S or N protein detection assays however they all suffered with very poor accuracy, reliability and sensitivity. Therefore it is a global consensus there is an urgent and tremendous demand for low-cost rapid and reliable SARS-CoV-2 antigen and virus detection which helps to enable timely and affordable point-of-care COVID-19 diagnostics for a very large population.

As the matter of fact, surface plasmon resonance (SPR) sensing is a promising label free and one-step virus detection method and has been used in rapid viral particle detection such as SARS/MERS^19^, H1N1^20^, H7N9^21^ Once the viral particles are captured by the monoclonal antibody immobilized on the SPR sensor chip surface, the plasmon resonance wavelength or intensity change induced by the virus particle presence can be measured by an optical sensing system^22-24^. However conventional SPR testing equipment are bulky and not affordable to most research and clinical institutions especially in developing countries and resources limited settings^25, 26^. Therefore the SPR viral detection, although often showed in research labs, rarely becomes a viable method accessible to clinical and point-of-care applications.

Here we report a low-cost nanoplasmonic sensor allowing for one-step rapid detection and quantification of the SARS-CoV-2 pseudovirus (Fig. 1a). The low-cost nanoplasmonic sensor chips were made by the method previously reported ^27,28^, and this method allows for large-scale manufacturing with high uniformity and repeatability (Fig. 1b). Owing to the specially designed periodic nanostructures, without any external coupling optics, the plasmon resonance wavelength and intensity change on the virus-capturing sensor can be simply observed by transmission light spectroscopy or imagin^29, 30^. Therefore the nanoplasmonic sensor chips can be integrated with microwell plate or microfluidic cuvette, and the measurements are carried out in both ubiquitous generic microplate readers and a low-cost handheld point-of-care testing device^27, 28^. The genetically reconstructed pseudovirus for SARS-CoV-2, SARS (Severe Acute Respiratory Syndrome), MERS (Middle East Respiratory Syndrome), and VSV (Vesicular Stomatitis Virus) viruses were produced for testing the device sensitivity and specificity^31-33^. With proper antibody functionalization on its surface, the nanoplasmonic sensor is able to detect as few as 30 SARS-CoV-2 pseudovirus while showing negligible responses to SARS, MERS and VSV pseudoviruses. SARS-CoV-2 virus concentrations in the range of 10^3^ vp/mL to 10^6^ vp/mL can also be quantified by this assay with simultaneous measurements of diluted standard samples in the same microplate reader. Similar sensing capability are demonstrated by using a low-cost handheld optical equipment controlled by a smartphone App. The ultrasensitive SARS-CoV-2 virus detection and potential early diagnosis of COVID-19 disease become available for point-of-care applications in clinics, roadside triage sites and even home settings.

**Fig. 1.**
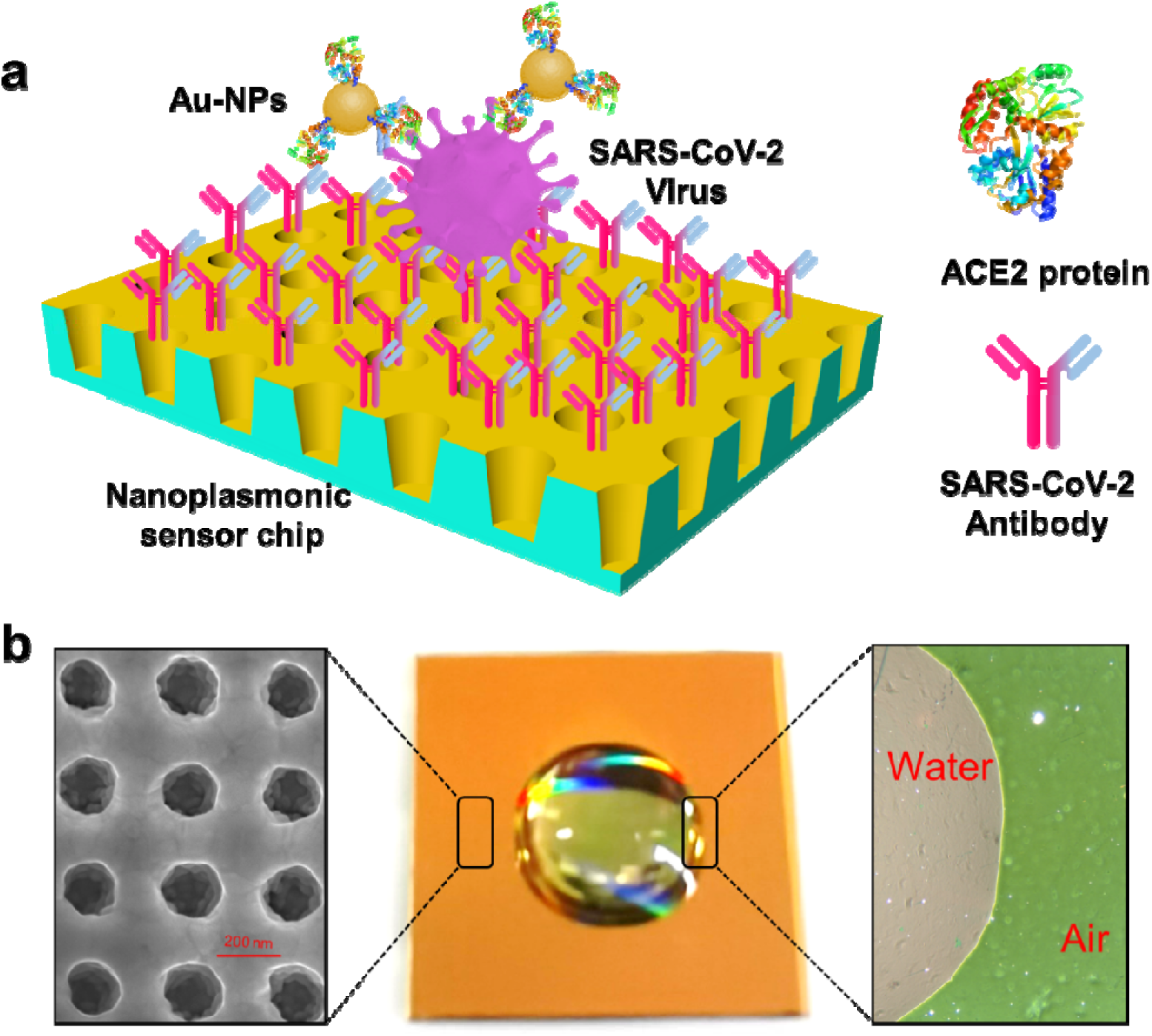
Label-free detection of SARS-CoV-2 pseudovirus with point-of-care devices. (a) Schematic diagram of nanoplasmonic resonance sensor for determination of SARS-CoV-2 pseudovirus concentration. (b) Photograph of one piece of Au-TiO2-Au nanocup array chip with a drop of water on top. Scanning electron microscopy (Left) shows the replicated nanocup array. Transmission microscopy image (Right). Air and water on the device surface show different colors, green and far red pink, respectively.

## Results

### Measurement of SARS-CoV-2 pseudovirus using the generic microplate readers with homemade 96-well chip plate device

Fast and sensitive detection of intact coronavirus directly in body fluid samples are essential to diagnose these viruses and help proper treatment. For this reason, our chip-in-microwell sensor were developed for the direct multichannel detection of different concentrations of whole virus particles rather than genetic extraction and analysis methods, which are accurate, but also time-consuming and low-cost (Fig. 2a,b). The surface of the nanoplasmonic array sensor chip was functionalized with SARS-CoV-2-specific antibodies to capture intact coronavirus by binding with spike proteins on the surface of the coronavirus (Fig. 2c). The initial titer after centrifugation of SARS-CoV-2 pseudovirus was 3.74 × 10^10^ vp/mL and it was quantified via the standard qRT-PCR method. Different dilutions of SARS-CoV-2 pseudovirus in PBS solution (from 1.0× 10^6^ to 1.6 × 10^10^ vp/mL) were added into the 96-well chip plate at the same time. The PBS solution in the absence of SARS-CoV-2 pseudovirus was treated as the control group. The real-time dynamic binding curves of the antigen-antibody interaction was shown in Fig. 3a. It can be seen that the curve of the control PBS solution group shows almost no change. Interestingly, the relative OD changes are proportional to these SARS-CoV-2 pseudovirus concentrations. The standard curve of the SARS-CoV-2 pseudovirus detection by the chip-in-microwell sensor over the range of 1.0× 10^6^ to 1.6 × 10^10^ vp/mL is shown in Fig. 3b. The relative OD change value was obtained from the data change over 60 min process as shown in Fig. 3a. The results were analyzed and fitted using a four-parameter logistic equation, and the correlation coefficient (R^2^) is 0.99246. Moreover, the R^2^ is 0.99994 in the relatively low virus concentrations range from the 1.0× 10^6^ to 2.5 × 10^8^ vp/mL.

**Fig. 2.**
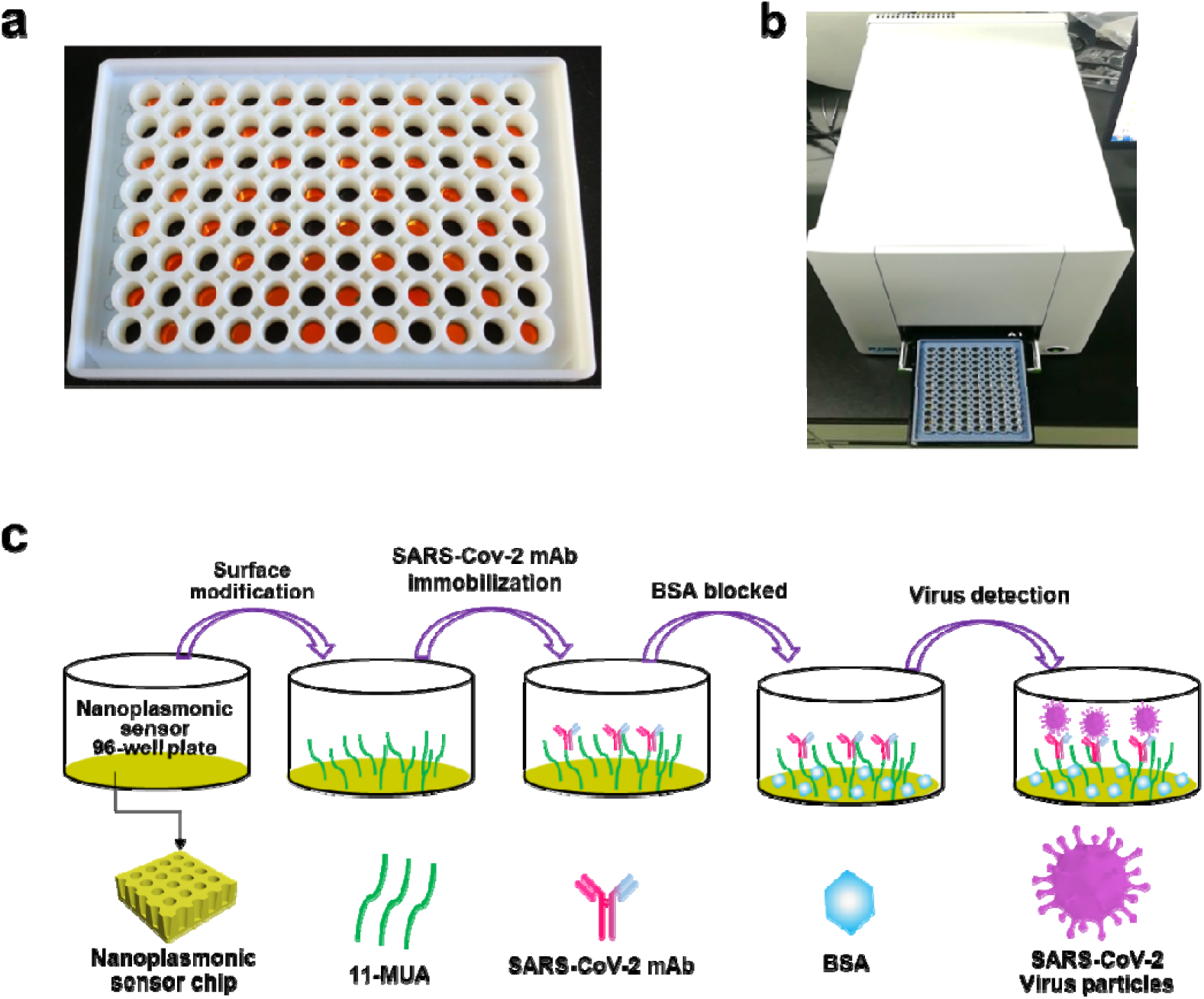
Nanoplasmonic sensor chip in microwell plate for detecting SARS-CoV-2 virus. (a) Integration of the nanoplasmonic sensor chip with a standard 96-well plate. (b) Testing with a generic microplate reader. (c) Schematic of nanoplasmonic sensor chip surface functionalization as well as capturing and detecting SARS-CoV-2 pseudovirus.

**Fig. 3.**
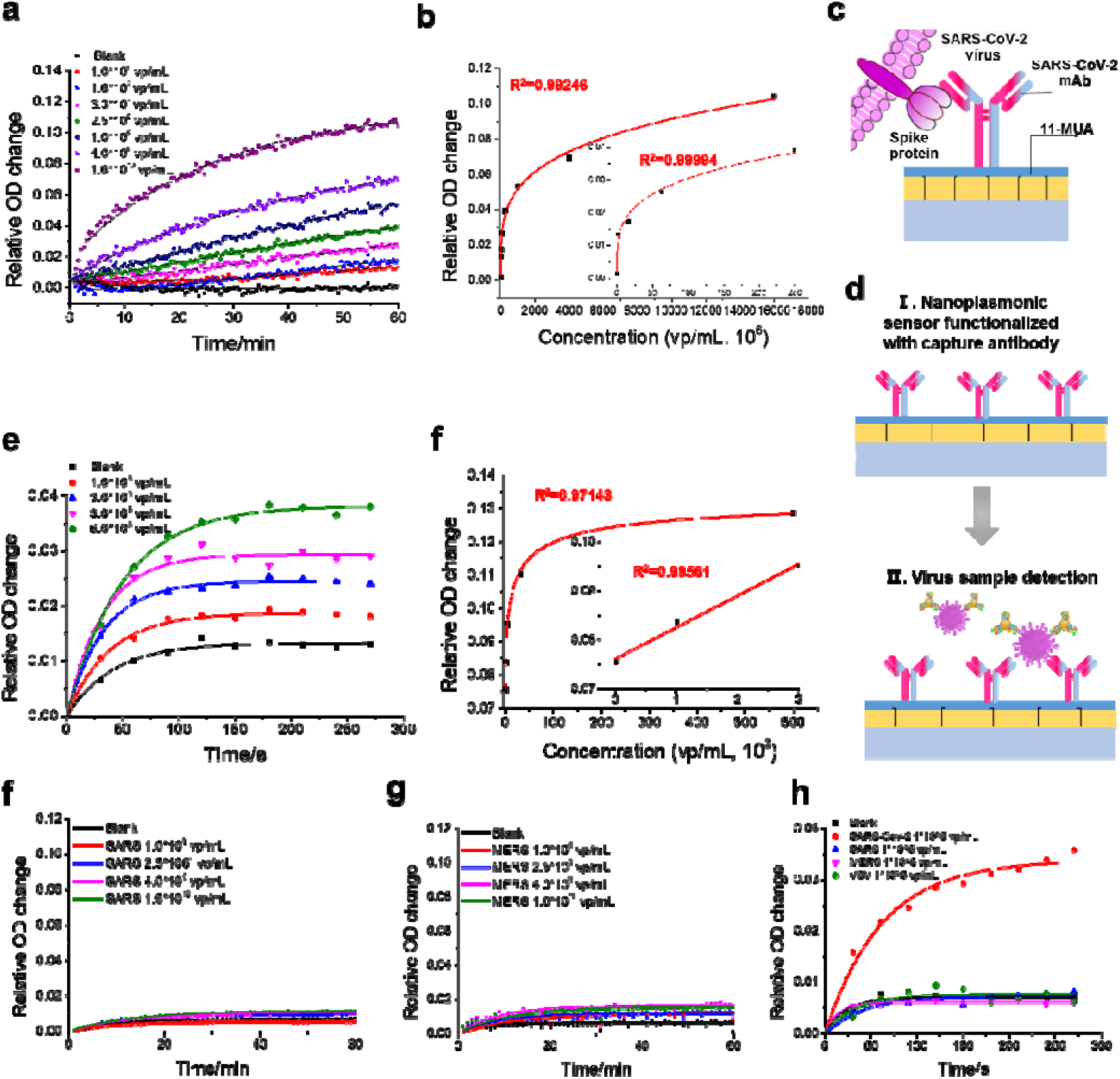
Detection of SARS-CoV-2 pseudovirus with nanoplasmonic sensor chip by a generic microplate reader. (a) Dynamic binding curves of SARS-CoV-2 antibodies interaction with different concentrations of the SARS-CoV-2 pseudovirus over the range 1.0× 10^6^ to 1.6 × 10^10^ vp/mL at the resonant wavelength. (b) SARS-CoV-2 pseudovirus standard curve (R^2^ = 0.99246, LOD = 30000 virus particles). (c) The illustration shows the binding of spike protein on the surface of SARS-CoV-2 virus to the specific SARS-CoV-2 mAbs. SARS-CoV-2 antibodies are conjugated to the activated carboxyl groups of 11-MUA on chip surface. (d) Schematic of nanoplasmonic sensor chip detecting SARS-CoV-2 pseudovirus with AuNP enhancement. (e) Au NPs-enhanced binding curves with different concentrations of the SARS-CoV-2 pseudovirus over the range 1.0 × 10^3^ to 6.0 × 10^5^ vp/mL in 30 uL samples. (f) Au NPs-enhanced SARS-CoV-2 pseudovirus standard curve (R^2^ = 0.97143, LOD = 30 virus particles). (f-g) Evaluation of the specificity of the nanoplasmonic sensor chip functionalized with SARS-CoV-2 antibodies. Dynamic binding curves of SARS-CoV-2 antibodies interaction with different concentrations of the SARS (f), and MERS (g). (h) pseudovirus were detected over the range 1.0× 10^6^ to 1.6 × 10^10^ vp/mL at the sensor resonance wavelength. (d) Specificity verification test of Au NPs-enhanced SARS-CoV-2 pseudovirus detection: Dynamic binding curves of SARS-CoV-2 antibodies interaction with different pseudovirus of SARS-CoV-2, SARS, MERS, and VSV at the concentration of 1.0× 10^5^ vp/mL.

Furthermore, we present a gold nanoparticles (NPs)-enhanced detection techniques based on our nanoplasmonic resonance sensor device that enables enhanced-sensitivity, rapid and convenience coronavirus detection in a single-step sandwich plasmonic resonance immunoassay method (Fig. 3c,d). We implemented the proposed techniques for detection of SARS-CoV-2 pseudovirus using the specific binding of SARS-CoV-2 mAbs and Au NP-labeled ACE2 protein to the SARS-CoV-2 spike protein and the receptor-binding domain (RBD) in SARS-CoV-2 spike protein, respectively. We firstly mixed 30 μL of different concentrations of SARS-CoV-2 pseudovirus with 10 μL of AuNP labeled ACE2 protein solution, and then dropped the mixture samples onto the spike protein specific nanoplasmonic resonance sensor for detection by using a generic microplate reader. The real-time nano-SPR curve of the interaction is represented in Fig. 3e. There is a clear distinction between different concentrations of SARS-CoV-2 pseudovirus within 5 min. Rapid quantitative detection of SARS-CoV-2 virus particles of a much lower concentration range(from 1.0× 10^3^ to 6.0 × 10^5^ vp/mL) was achieved via a single-step Au NP-enhanced nanoplasmonic detection techniques based on our homemade 96-well biosensor chip plate device. In addition, the 4PL curve fitting R^2^ between the relative OD change value and the concentrations of SARS-CoV-2 pseudovirus is 0.97143 over the range of 1.0× 10^3^ to 6.0 × 10^5^ vp/mL, and the R^2^ is improved up to 0.99561 in the lower concentration range from 1.0× 10^3^ to 3.0 × 10^3^ vp/mL (Fig. 2f). The limit of detection (LOD) of the chip-in-microwell biosensor for the SARS-CoV-2 pseudovirus detection was estimated to be 30 virus particles in the final sample solution. Notably, this LOD is well below the typical viral concentration range in nasopharyngeal swab and saliva^3, 34, 35^, suggesting that the chip-in-microwell sensor has the potential to detect SARS-CoV-2 virus with an ultrahigh sensitivity and effectiveness in early infection diagnostics, compared to existing technologies requiring laborious sample processing and amplification.

Reliable detection of SARS-CoV-2 virus requires distinguishing nonspecific binding of other viruses to the functionalized nanoplasmonic sensor surface. Virus selectivity is achieved by surface immobilized highly specific antibodies CR3022 showing strong affinity only to the SARS-CoV-2 coronavirus membrane proteins^1^, called spike proteins (Fig. 3c). Here, the detection specificity was evaluated with the SARS, MERS, and VSV in comparison with SARS-CoV-2 pseudovirus. As shown in Fig. 3f-h, a significant difference in binding capacity was observed between the SARS-CoV-2 viruses and other virus strains. These results demonstrate that the functionalized nanoplasmonic sensor chip has very high specificity (>1000:1) in detecting the SARS-CoV-2 virus.

### Measurement of SARS-CoV-2 pseudovirus using a low-cost handheld optical equipment controlled by a smartphone App

Demand for rapid, accurate and convenient SARS-CoV-2 virus detection present significant challenges in controlling and stopping the pandemics. Diagnosis of patients in the early stage infection are so far limited to viral nucleic acid or antigen detection in human nasopharyngeal swabs or saliva samples. Although traditional approaches, including point-of-care (POC) diagnostics, bedside testing, and community-based approaches, were applied to address these challenges, innovative techniques combining with mobile technologies, nanotechnology, imaging systems, and microfluidic technologies are expected to promote this transformation^36-38^. In this work, we also developed a portable and innovative devices controlled by a smartphone App for real-time measurements of the dynamic binging curves of SARS-CoV-2 virus on the nanoplasmonic sensor (Fig. 4a). We integrated the nanoplasmonic sensor chip in a cartridge designed for the handheld testing device, followed by functional modification of the sensor chip and detection of pseudovirus particle samples according to the protocol described previously (Fig. 4b). The functionalized chip cartridge with different concentrations of pseudovirus samples was inserted into the testing device and the dynamic curves were recorded in real time through the smartphone APP. The real-time virus binding curve measurement is presented in Fig. 4c and Video 1. This low-cost handheld sensing platform can directly detect the SARS-CoV-2 pseudovirus sample in one step within 15 minutes and the detectable virus concentrations range over 8.0 × 10^4^ to 6.0 × 10^6^ vp/mL. Moreover, the detection limit of the handheld equipment is currently about 4000 SARS-CoV-2 virus particles and can be further improved to be comparable with the microplate reader case.

**Fig. 4.**
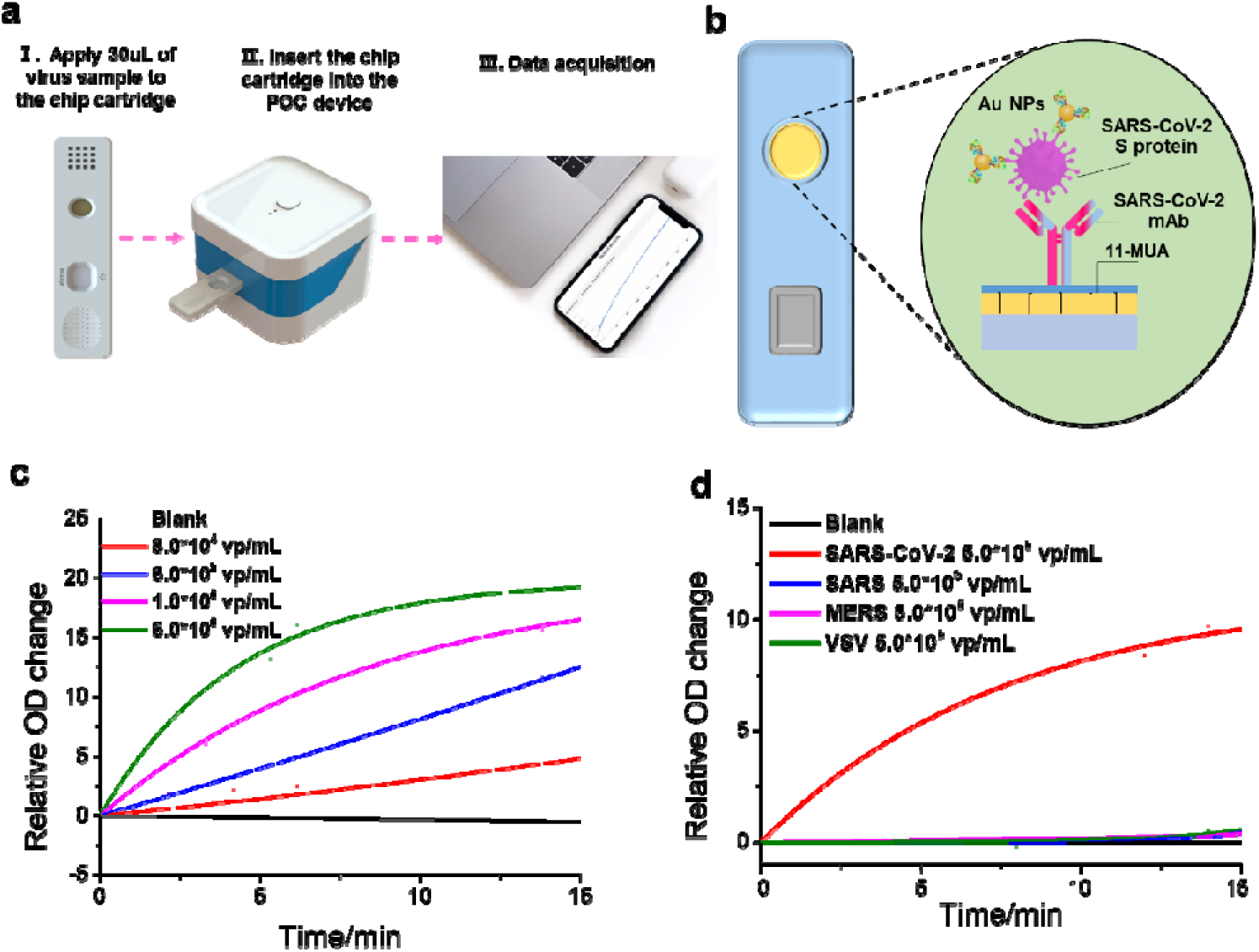
Detection of SARS-CoV-2 pseudovirus with nanoplasmonic sensor chips by a point-of-care device. (a) Schematic of nanoplasmonic sensor chip cartridge detecting SARS-CoV-2 pseudovirus with a low-cost handheld point-of-care testing device. (b) The illustration shows the detection process of the sensor chip cartridge for specific SARS-CoV-2 detection. (c) Dynamic binding curves of virus and antibody interaction with different concentrations of the SARS-CoV-2 pseudovirus over the range 8.0× 10^4^ to 6.0 × 10^6^ vp/mL at the resonance wavelength. (d) Specificity verification test: Dynamic binding curves of SARS-CoV-2 antibodies interaction with different pseudovirus of SARS-CoV-2, SARS, MERS, and VSV at the concentration of 5.0× 10^5^ vp/mL.

Furthermore, the detection specificity of the handheld devices for SARS-CoV-2 pseudovirus was also carried out with the SARS, MERS, and VSV pseudovirus. As shown in Fig. 4d, there was no obvious change in the curves of SARS, MERS, and VSV pseudovirus. A remarkably rising curve was observed only for the SARS-CoV-2 pseudovirus sample, suggesting that the devices controlled by a smartphone App could offer convenient operability while permit highly sensitive and specific quantification and detection of SARS-CoV-2 pseudovirus. This nanoplasmonic sensor device with the potential in rapid and affordable early diagnosis of COVID-19 disease will become available for POC applications in clinics, roadside triage site and even home settings.

## Conslusion

At present the COVID-19 pandemic is still affecting the whole world. However, there are just a limited rapid diagnostic methods or testing equipments that are effective for newly infected patients or asymptomatic carriers. In addition, most detection methods for SARS-CoV-2 viruses have high logistical barriers and thus are not suitable for POC testing. Therefore in this work, we developed a simple and low-cost device to rapidly and sensitively detect the SARS-CoV-2 virus in one step using a nanoplasmonic biosensor integrated with a standard 96-well plate. Our study confirmed that the nanoplasmonic sensor chip is able to directly detect the whole SARS-CoV-2 virus with extraordinary time efficiency (<15 min) and sensitivity (LOD = 30 virus particles). Moreover, similar rapid and sensitive detection capabilities are demonstrated by using a low-cost handheld optical equipment controlled by a smartphone App. The detection limit of SARS-CoV-2 pseudovirus in the final detection system was about 4000 virus particles within 20 min. We therefore conclude that the low-cost POCT devices would be expected for rapid diagnosis of SARS-CoV-2 virus infection.

## Experimental Section

### Materials

Hexylsilane, 6-mercapto-1-hexanol (MCH), 11-mercaptuoudecanoic acid (MUA), 1-ethyl-3-(dimethylaminopropyl) carbodiimide (EDC), N-hydroxysuccinimide (NHS), bovine serum albumin (BSA), ethanolamine and phosphate-buffered saline (PBS) buffer were purchased from Sigma-Aldrich, SARS-COV-2 Antibody-CR3022 (catalog no. CHA005) was purchased from Sanyou Biopharmaceuticals Co., Ltd. Angiotensin-Converting Enzyme 2 (ACE2) Protein (catalog no. 2020T13) was purchased from Shanghai EasyBiotech Co., Ltd.. All chemicals were used as obtained without any further purification. The SARS-CoV-2 pseudovirus samples were obtained from Shanghai public health clinical center.

### Fabrication process

The nanoplasmonic sensor was fabricated by a replica molding process and parameters of sensor chip are described in Supporting Information. The original mold was a tapered nanopillar array on silicon wafer made by lithography and plasma etching. An optical adhesive liquid was evenly spread on the mold and placed on a polyethylene terephthalate (PET) sheet. After 3 min of UV light (105 mW/cm^2^) irradiation, the PET sheet with the UV polymer layer was peeled off from the mold. Then, 10 nm of titanium (Ti) and 70 nm of gold (Au) were deposited onto the nanocup array in an electron beam evaporator. Later the sheet were cut into small pieces of 1 cm×1 cm, and glued to an open-bottom 96-well plate or a chip cartridge made by a 3D printer (Object 30 primer(tm) Stratasys Ltd.).

### Surface functionalization

After rinsing the chip integrated microplate wells or cards twice with deionized (DI) water, the chips were incubated in a 10 mM MUA ethanol solution for 0.5 h at room temperature. Clean the chips with 70% ethanol and DI water twice followed by immersion in a mixture of 400 mM EDC and 100 mM NHS for 30 min to activate the chips at room temperature. Next, the chips were rinsed in DI water twice and the SARS-CoV-2 mAbs of 12.0 μg/mL (in PBS) were immediately immobilized to the activated chips for 4 h at room temperature. Rinse the chips with PBS and DI water, and afterwards incubate the chips in 60 μg/mL BSA blocking solution and then in 10% ethanolamine solution, and both steps last for 30 min at room temperature. After incubation, functionalized chips were gently rinsed with DI water twice and then stored in a humid atmosphere at 4 °C.

### Preparation of gold labeled ACE2 protein

ACE2 Protein prepared in pH 7.0 Glycine-buffered solution (100 nmol/L) was added drop-wise to 1 ml of 30 nm colloidal Au-NPs solution. The pH of the colloidal Au-NPs solution was adjusted to pH 7.2 by addition of dilute 0.01 M K_2_CO_3_ before adding the ACE2 protein. This was done in as much as the optimum stability of the conjugates is at pH 7.2, and the least content of antibody was 0.023 mg/ml of gold solution. The ACE2 protein was initially incubated with Au-NPs for 30 min. After incubation, 10 μL of Tris-HCl 1× with 1% BSA blocking solution was added to the gold labeled solution for 10 min. The Au-NPs suspension was then centrifuged at 8000 rpm for 30 min. Centrifugation and resuspension steps were repeated 3 times. At the final resuspension step, Au-NPs precipitate were resuspended in 100 μL of Tris-HCl 1× with 0.2% w/v PEG and 0.05% w/v Tween20. The conjugated Au NPs were stored at 4 °C for further experiments.

### Measurement of SARS-CoV-2 pseudovirus using the homemade 96-well chip plate device

The dynamic interaction kinetics of the SARS-CoV-2 pseudovirus particles binding to the immobilized SARS-CoV-2 mAbs with respect to time was performed using the generic microplate reader with our homemade chip-in-microwell plate. 100 μL of different concentrations of pseudovirus particles (covering the range of 10^6^ vp/mL to 10^10^ vp/mL) were added into the functionalized chip-in-microwell plate at the same time. The dynamic absorption spectrum corves were measured using a simple small volume microplate reader (Xlement SPR100). The Au NP-enhanced techniques for detection of SARS-CoV-2 pseudovirus was implemented by using the specific binding of SARS-CoV-2 mAbs and Au NP-labeled ACE2 protein to the SARS-CoV-2 spike protein and the receptor-binding domain (RBD) in SARS-CoV-2 spike protein, respectively. We firstly mixed 30 μL of different concentrations of SARS-CoV-2 pseudovirus with 10 μL of gold labeled ACE2 protein solution, which was followed by dropping the mixture samples into the spike protein specific nanoplasmonic resonance sensor for detection via the generic microplate reader.

## Data availability

All relevant data are available from the authors.

## Acknowledgments

This work was supported by National Natural Science Foundation of China (91959107),Huazhong University of Science and Technology (2020kfyXGYJ111). The authors thank Lu Wang, Hong Yang and Hui Guo for their help with data acquisition.

## Author contributions

**Liping Huang**: Conceptualization, Methodology, Investigation, Data curation, Visualization, Writing - original draft. **Longfei Ding**: Pseudovirus production and quantification. **Jun Zhou**: Chip manufacturing. **Shuiliang Chen**: Chip manufacturing. **Fang Chen:** Chip manufacturing. **Chen Zhao:** Pseudovirus design and antibody selection. **Yiyi Zhang**:Software. **Wenjun Hu**: Investigation, Resources. **Jiansong Ji:** Medical advisory. **Hao Xu**: Supervision, Funding acquisition. **Gang L. Liu**: Supervision, Writing - review & editing, Project administration.

## Additional information

Competing financial interests: The authors declare no competing financial interests.

